# Gene expression signatures of response to fluoxetine treatment: systematic review and meta-analyses

**DOI:** 10.1101/2024.02.19.581045

**Authors:** David G. Cooper, J. Paige Cowden, Parker A. Stanley, Jack T. Karbowski, Victoria S. Gaertig, Caiden J. Lukan, Patrick M. Vo, Ariel D. Worthington, Caleb A. Class

**Author notes:** Corresponding Author: Caleb A. Class, 4600 Sunset Ave., Indianapolis, IN, USA, 46208.

## Abstract

**Background:** Selecting the best antidepressant for a patient with major depressive disorder (MDD) remains a challenge, and some have turned to genomic (and other ‘omic) data to identify an optimal therapy. In this work, we synthesized gene expression data for fluoxetine treatment in both human patients and rodent models, to better understand biological pathways affected by treatment, as well as those that may distinguish clinical or behavioral response.

**Methods:** Following the PRISMA guidelines, we searched the Gene Expression Omnibus (GEO) for studies profiling humans or rodent models with treatment of the antidepressant fluoxetine, excluding those not done in the context of depression or anxiety, in an irrelevant tissue type, or with fewer than three samples per group. Included studies were systematically reanalyzed by differential expression analysis and Gene Set Enrichment Analysis (GSEA). Individual pathway and gene statistics were synthesized across studies by three p-value combination methods, and then corrected for false discovery.

**Results:** Of the 74 data sets that were screened, 20 were included: 18 in rodents, and two in tissue from human patients. Studies were highly heterogeneous in the comparisons of both treated vs. control samples and responders vs. non-responders, with 737 and 356 pathways, respectively, identified as significantly different between groups in at least one study. However, 19 pathways were identified as consistently different in responders vs. non-responders, including toll-like receptor (TLR) and other immune pathways. Signal transduction pathways were identified as consistently affected by fluoxetine treatment in depressed patients and rodent models.

**Discussion:** These meta-analyses confirm known pathways and provide new hints toward antidepressant resistance, but more work is needed. Most included studies involved rodent models, and both patient studies had small cohorts. Additional large-cohort studies applying additional ‘omics technologies are necessary to understand the intricacies and heterogeneity of antidepressant response.

## INTRODUCTION

Despite decades of advances in the treatment of depression, approximately half of patients do not respond to their first prescribed antidepressant(1,2). Nonresponsive patients may spend years cycling through the available therapies before finding one that works; some will remain treatment-resistant, which is associated with higher risk of suicide(3). Advancements in precision medicine have led to modest improvements in treatment response; for example, one clinical trial demonstrated a 5% improvement in response vs. treatment as usual when the antidepressant was prescribed based on a genetic test involving 59 alleles and variants across eight genes(4–6). Meanwhile, many novel or repurposed drugs are under investigation or in trials as alternative or adjunct therapies for depression; while this will provide additional options for those suffering from treatment-resistant depression (TRD), selecting from the increasing number of approved antidepressants will remain challenging(5).

Selective serotonin reuptake inhibitors (SSRIs) are currently the most commonly prescribed antidepressant class, in large part due to relatively high tolerability and efficacy(7,8). As the first SSRI to win regulatory approval in the United States, and one of the best tolerated, fluoxetine remains one of the most prescribed antidepressants, and many research studies have been conducted to understand its therapeutic efficacy and mechanism of action(9,10). Use of computational chemistry and “omics” methods have increased scientific knowledge in this and many other areas, providing leads for new drugs, better insights into their mechanisms of action, and potential signatures of response(5,11,12). However, these methods have not been the panaceas that some may have hoped, as heterogeneity between patients, interactions between multiple levels of biology, and environmental and other factors can be difficult to capture(2). So while progress has been made understanding the mechanism of action of antidepressants, exact causes of differences in response between patients remain enigmatic.

The use of systematic reviews in mental disorders and other fields has dramatically increased in recent years to synthesize the wealth of clinical data being generated(13–16). However, these guidelines have been applied less frequently in the use of gene expression data, including one non-PRISMA systematic review and meta-analysis of gene expression signatures corresponding to fluoxetine treatment in rodents(17–19). These studies generally focus on a specific tissue type in either humans or rodents, and translating between organisms remains difficult despite recent advances. Although over 15,000 annotated human genes have orthologs in mice, their function is not always shared, which has pharmacological implications; in cancer for example, fewer than 8% of drugs successfully translate from animal models to clinical trials(20–22). Niculescu *et al.* have employed convergent functional genomics to identify consistent genes in independent studies of patients and rodent models for risk prediction in depression, schizophrenia, and other psychiatric disorders to find biomarkers of clinical value(23–25). Alternatively, meta-analyses considering biological pathways do not find specific biomarkers but allow us to focus on the shared biological effects between studies, which may be more consistent across organisms(26,27). In this systematic review and meta-analysis, we will apply this approach to summarize and potentially identify new biological pathways of antidepressant treatment and response.

The objectives of this systematic review are to synthesize the evidence for gene expression modification by fluoxetine treatment, and whether gene expression levels distinguish clinical or behavioral response to treatment, across multiple tissue types in humans and rodent models. In this paper, we use “Response Signatures” to refer to gene expression distinguishing those with good vs. poor response to fluoxetine, while “Treatment Signatures” signifies differences between fluoxetine treatment vs. control. To our knowledge, this is the first systematic meta-analysis of gene expression data to investigate behavioral or clinical response to an antidepressant, as well as the first to integrate biological pathways across multiple organisms and tissue types. We apply a consistent analysis pipeline across all studies, so that we do not rely on varying definitions of statistical significance applied by different researchers. Results of the meta-analyses may be applied to improving the prediction and assessment of fluoxetine response shortly after treatment, and they may provide hints for combination therapies or new drug development for TRD.

## METHODS

The Gene Expression Omnibus (GEO) was identified as the main database for this systematic review, due to its primary focus on gene expression data and the relative consistency of data deposits and formatting(28). PubMed was identified as a contingent database if the GEO search resulted in fewer than five studies that passed screening. As one of the most prescribed and studied antidepressants, fluoxetine was selected as the focus of this systematic review; we were more inclusive regarding organisms and tissue type, allowing us to identify both heterogeneity and potential consistencies. The GEO search was conducted on 4 April 2023, using the following keywords: *(fluoxetine) OR (selective serotonin reuptake inhibitor) OR (ssri)*. Results were filtered within GEO using *Entry type = Series* (to return full data sets rather than individual samples), *Organism = homo sapiens, mus musculus,* or *rattus norvegicus*, and *Study type = Expression profiling by array* or *expression profiling by high throughput sequencing*. The resulting data series were manually filtered by authors DC and CC based on the exclusion criteria: studies were excluded if they were not primarily focused on depression or anxiety, were not conducted in a relevant tissue type or genetic background (cancer cell lines, for example), did not involve fluoxetine treatment, or if there were fewer than three samples per group. Decisions were made based on review of the GEO series abstract and sample metadata. If unclear, any cited publication and supplementary materials were consulted for additional information. For synthesis, studies would be grouped based on the comparison: good vs. poor response, fluoxetine treatment vs. control, or fluoxetine treatment vs. control (within depressed patients and rodent models, and separately within unstressed rodents). Studies would be synthesized both across and within organisms.

Data extraction, transformation, synthesis, and assessment were conducted using R version 4.2.3(29). Each individual analysis was completed by one author and checked by DC or CC. All analysis code and results are available for download (see *Code Availability*). Gene expression data were downloaded using the R library GEOquery, and then checked for completeness(30). Data sets that were not retrieved properly using GEOquery were downloaded directly from the GEO website and imported into R for analysis. Basic quality control was assessed by visualizing and comparing gene expression distributions, principal components analysis, and between-sample correlations. Clear outlier samples were removed prior to any subsequent analyses. Gene-level differential expression analysis was conducted using the R libraries DESeq2 for RNA sequencing data, or limma for microarray data(31,32). All differential expression analyses were conducted as *group* vs. *group*, with the main variables being treatment status or response (as appropriate). Other variables sought were tissue type, time point, and participant/rodent ID. For each comparison, the effect measures used in subsequent steps were the log2 fold change (log2FC), nominal p-value, and t-statistic. When multiple tissue types were present in a study, separate analyses were conducted within each tissue. Risk of bias was assessed by evaluating the provided characteristics of the study populations.

Differential expression results for each comparison were summarized to the pathway level using the fgsea R library for Gene Set Enrichment Analysis (GSEA), using the Reactome and KEGG pathway gene sets accessed through the Consensus Pathway Database (CPDB)(33–37). The CPDB provides orthologous gene sets for human and mouse genomes, allowing for the synthesis of human and rodent models at the pathway level (for rat data sets, gene ID’s were converted to mouse ID’s by homology prior to GSEA). A ranked list of differential expression t-statistics from each comparison was input to fgsea, and relevant effect measures for each pathway were the normalized enrichment score (NES) providing the magnitude of gene expression enrichment in one group or the other, and nominal p-value.

Results across comparisons were synthesized using MetaDE(38). P-value combination methods were selected due to substantial heterogeneity between studies (organism, tissue type, and gene expression platform). Within organisms, p-values were synthesized at both the gene and pathway level, while across-organism syntheses were only conducted at the pathway level. Fisher’s method was used to identify genes or pathways that were differentially expressed in *any study*, while Wilkinson’s method (*Max-P*) was used to identify genes or pathways that were consistently differentially expressed *across studies*(39–41). The resulting meta-analysis p-values were corrected for false discovery using the method of Benjamini and Hochberg to result in q-values(42). As an intermediate method, we also selected genes or pathways with nominal p<0.05 in greater than half of synthesized comparisons (Frequency of 50%, or *Freq50*). A depiction of these methods is provided in **Figure 2A**. Synthesis results were displayed using Venn diagrams to show how many genes/pathways were identified by each meta-analysis method, and bar graphs showing pathways identified by Max-P with q<0.05.

**Figure 1.**
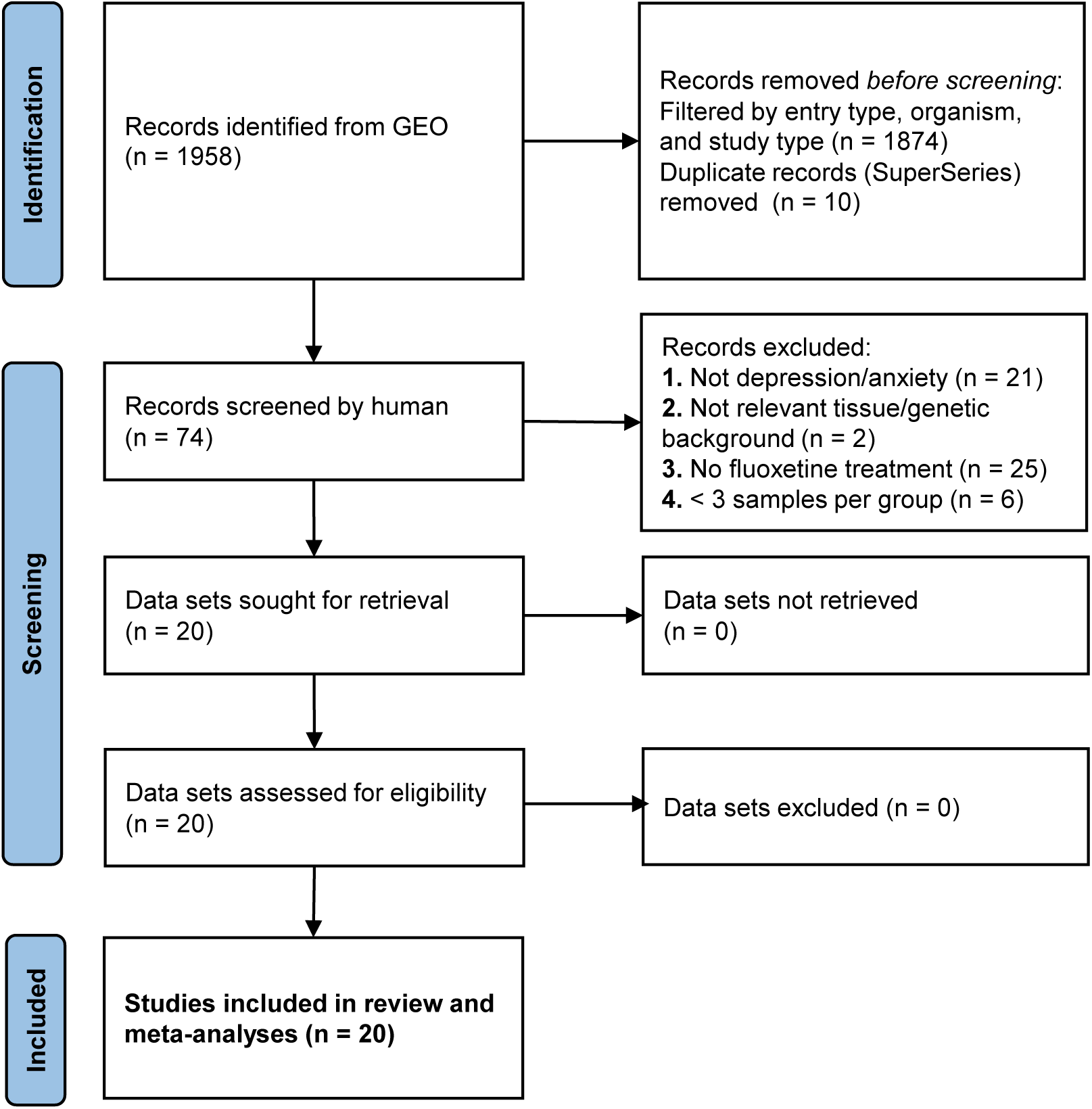
PRISMA flow diagram of systematic search and screening for gene expression studies assessing fluoxetine treatment.

**Figure 2.**
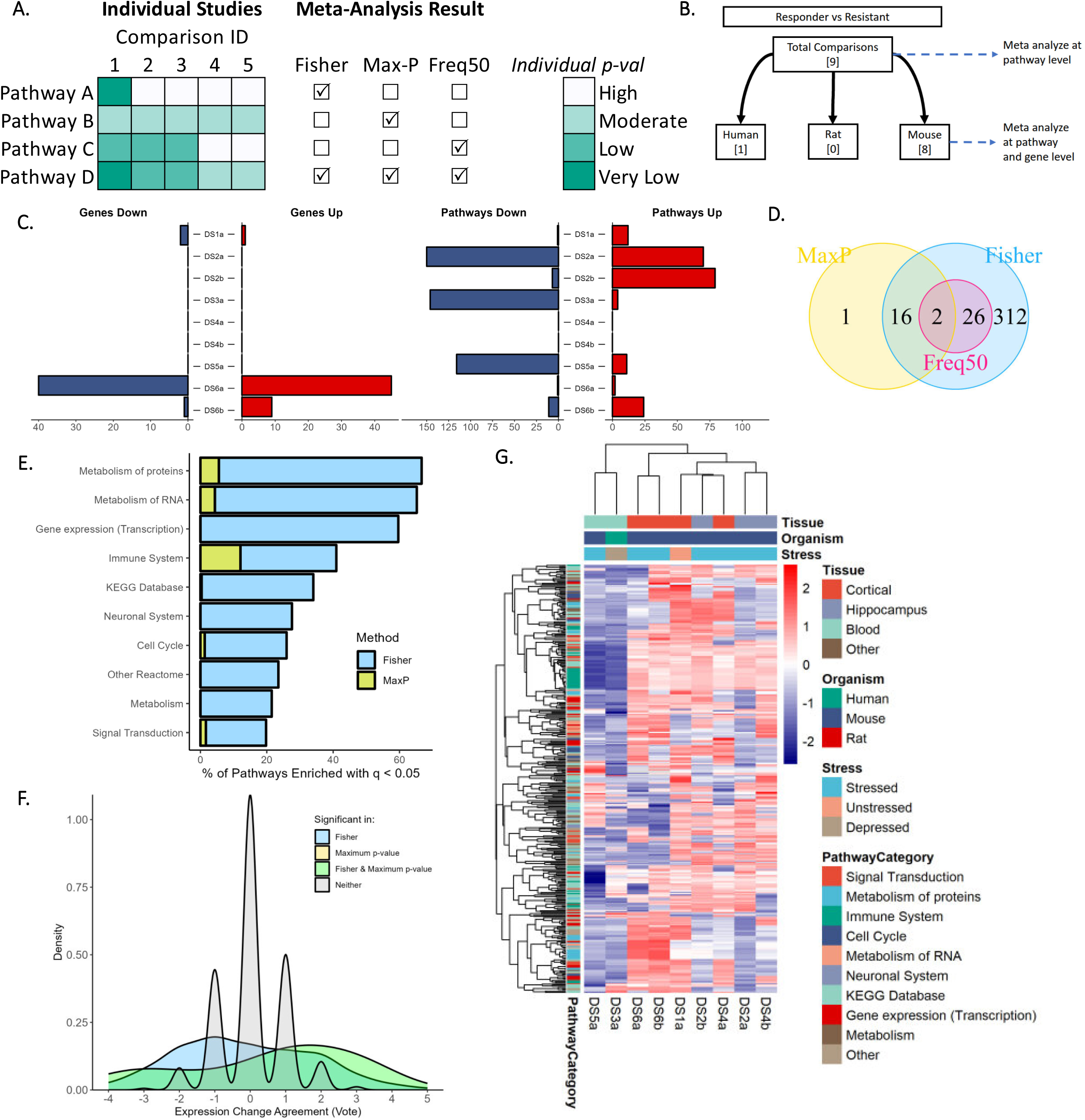
(A) Demonstration of meta-analysis methods used in this study to synthesize pathway analyses and differential expression analyses. Heatmap on left shows GSEA nominal p-values for four example pathways in five comparisons. Checkboxes indicate which meta-analysis method(s) would identify that pathway as significantly enriched. **(B)** Individual comparisons used in meta-analysis for gene expression signatures of behavioral or clinical response. **(C)** Number of genes differentially expressed (left) and pathways enriched (right) in each of the nine comparisons of samples from responders vs. non-responders (q < 0.05). “Up” indicates greater expression in responders. **(D)** Pathways identified as significantly enriched (q < 0.05) across the nine comparisons by the three meta-analysis methods. **(E)** Percentage of pathways significantly enriched in responders or non-responders from each of the Reactome categories (“Top Level”) or KEGG Database. x-axis indicates percentage of pathways from a certain category identified as enriched. **(F)** Density plot showing distribution of Vote Sums across pathways, colored by meta-analysis result. Positive scores indicate enrichment in responders, negative in non-responders **(G)** Heatmap of Normalized Enrichment Score (NES) across comparisons for pathways identified as enriched in any comparison (Fisher’s q < 0.05). Positive NES indicates enrichment in responders.

Heterogeneity among study results was investigated using subgroup analysis within organisms or tissue types, as well as heatmaps of pathway NESs across studies. To estimate certainty of the meta-analyses, consistency of the direction of enrichment was estimated by simple vote-counting across comparisons: a pathway or gene was assigned +1 for each comparison where it was significantly enriched in responders with nominal p<0.05, or -1 if enriched in non-responders. These were summed across all comparisons, and numbers further from zero indicate greater consistency in results that are statistically significant within comparisons. For genes or pathways identified as consistently differentially expressed across studies, we generally expect them to agree in the direction of the effect (i.e. overexpressed in responders or non-responders). This is not assured, as differing and even opposite effects have been reported in some cases(43,44).

Sensitivity analyses were performed to assess the meta-analyses. Of nine response vs. non-response comparisons, eight involved mice while the other included human patients, so we looked at inclusion/exclusion of the patient data set. We further explored the effect of tissue type by removing the single comparison profiling blood in mice, resulting in seven comparisons in brain tissue only. Of the 13 treated vs. untreated experiments in unstressed mouse models, 12 involved microarray profiling in male mice, while the other used RNA-Seq in female mice; we conducted sensitivity analysis to investigate the effects of inclusion/exclusion of this last study in meta-analysis. Finally, we compared meta-analysis results of treatment signatures when DS19 (containing 27 comparisons from various brain regions) was removed.

## RESULTS

The initial keyword search resulted in 1958 entries, which was filtered to 84 data sets (known in GEO as “series”) using automatic filtering for Entry Type, Study Type, and Organism (**Figure 1**). Most of the automatically excluded entries were individual samples, which were not targets for this study and generally were included in one of the returned data sets. 10 “SuperSeries” were removed as duplicates, as they contained one or more individual series returned by the search. 74 data series were manually assessed for the exclusion criteria (**Supplementary Table 1**), resulting in 20 selected for inclusion(45–60) (**Table 1**, **Figure 1**). Of the 20 included data sets, two profiled tissue from patients diagnosed with Major Depressive Disorder (MDD): one profiled gene expression in whole blood from adolescent females before and after eight weeks of continuous fluoxetine treatment(57), while the other profiled lymphoblastoid cell lines (LCLs) developed from 10 patients with known antidepressant response based on change in score by Hamilton Depression Rating Scale (HDRS). These LCLs were treated with fluoxetine or control *in vitro* for three weeks(47). Six of the 10 patients were classified as responders, while the other four were classified as non-responders.

**Table 1.**
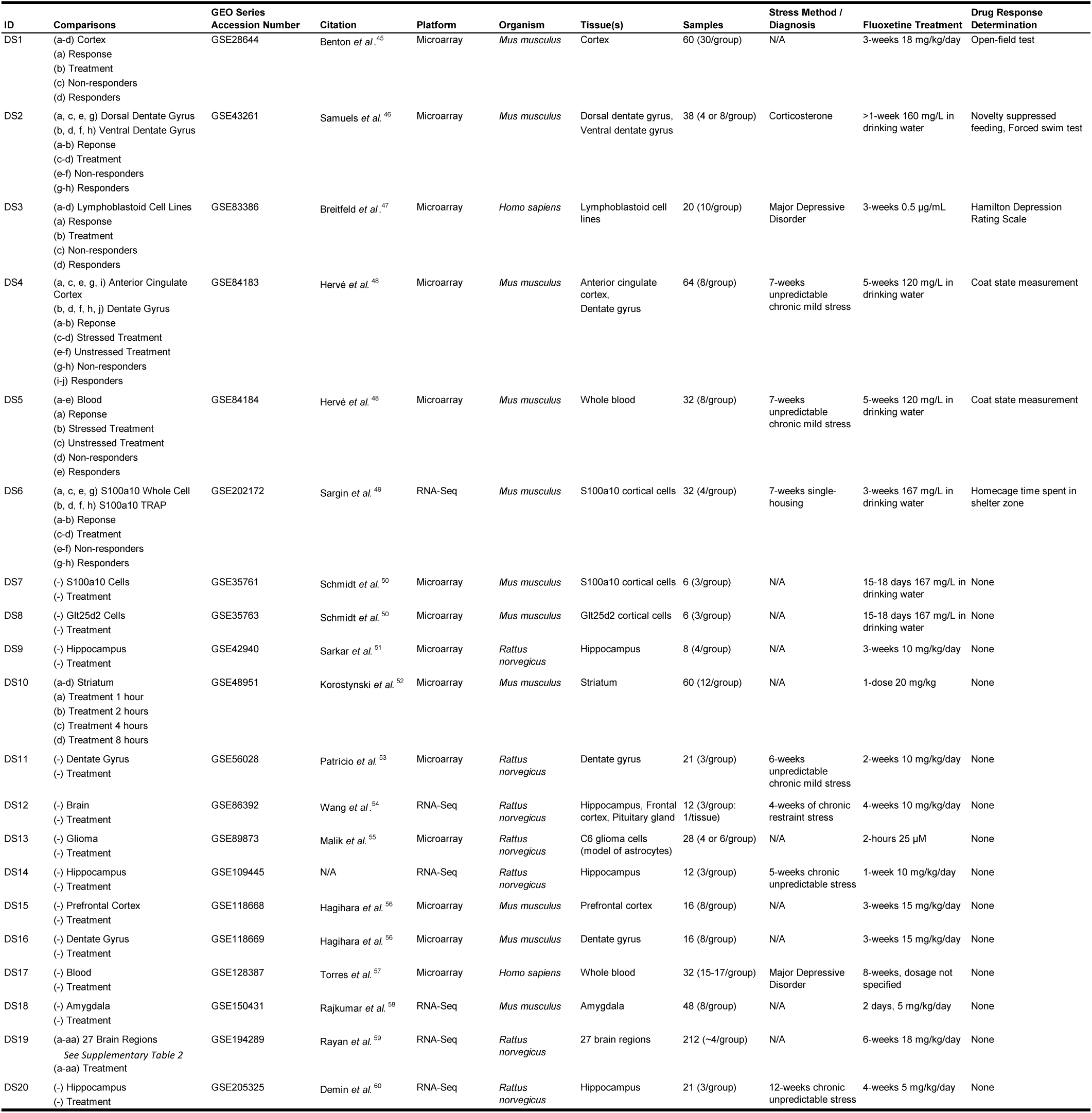
20 data sets included in synthesis. Letters in parentheses in “Comparisons” column are used as reference in subsequent figures.

The remaining 18 studies profiled gene expression with fluoxetine treatment in rodent models (11 in mice, seven in rats). Stress models were applied in nine of these studies to achieve anxious and/or depressive behavior: five employed chronic mild stress(48,53,60–62), while the others used either chronic restraint stress(54), single housing(49), injection with corticosterone(46), or selective breeding for high anxiety(55). Five studies included a classification of response in the treated animals, which was assessed using a behavioral method such as the open-field test (OFT) for measurement of anxiety(45) or the forced swim test (FST) for despair(46). One study did not induce stress but did measure behavioral response: anxiety was measured by OFT after three weeks of fluoxetine or vehicle treatment, and response was defined based on the ratio of scores between the fluoxetine and vehicle groups(45). One mouse study profiled whole blood, but the majority collected samples from one or more brain regions for gene expression profiling: the most common was hippocampal tissue (particularly dentate gyrus, included in five studies(46,48,53,56,59)). One profiled fluoxetine effects across 27 brain regions in rats, identifying region-specific differential expression signatures in both bulk and single-cell RNA-Seq data (we did not include single-cell data in this work) (**Supplementary Table 2**)(59).

As is common with rodent models, males were utilized in 17 of the 18 studies, demonstrating a substantial risk of bias. One study profiled gene expression with fluoxetine and imipramine treatment in female mice, which had been shown to exhibit behavioral despair by FST and tail suspension test (TST) after reduction in *Brd1* expression (this was not observed in male mice with reduced *Brd1* expression)(58). Conversely, the two human studies were biased toward female inclusion, with one including adolescent females only(57), and the other including eight females and two males in the microarray analysis(47). For the six data sets synthesized for Response Signatures, all five mouse data sets included males, while the patient cohort was biased toward females.

### Response Signatures

Meta-analysis by Fisher’s method, the Max-P method, and the Freq50 method (**Figure 2A**) were applied on nine comparisons from six data series to synthesize gene expression signatures associated with clinical or behavioral response to fluoxetine (**Figure 2B**). Systematic re-analysis of individual comparisons resulted in widely varying numbers of differentially expressed genes and pathways after false discovery correction for multiple testing (**Figure 2C**). DS6a had the greatest number of significant genes at 85, while four comparisons showed no statistically significant differential expression among genes with q<0.05. Conversely, statistically significant pathway enrichment was observed for all studies except DS4. Nominal p-values were synthesized across all comparisons for each Reactome and KEGG pathway and then corrected for false discovery to result in q-values. After false discovery correction, meta-analysis identified 356 pathways (over 30% of the pathways analyzed) enriched in at least one comparison (q<0.05 by Fisher’s method), 19 pathways enriched across comparisons (Max-P), and 28 pathways enriched in at least five comparisons (Freq50) (**Figure 2D**). Pathways involving metabolism of proteins or RNA, transcription, or the immune system were most likely to be enriched (**Figure 2E**).

We then considered direction of enrichment through vote-counting across comparisons: a pathway was assigned +1 if significantly enriched in responders, or -1 if enriched in non-responders. The distribution of vote sums for all pathways is provided in **Figure 2F**. Pathways that were not identified as enriched by either meta-analysis method had distributions centered around zero, with 92% of pathways ranging from -1 to +1, indicating the expected low agreement. Those identified as significant by meta-analysis had a wider range, indicating more consistent enrichment in responders or non-responders. However, the greatest vote sum magnitude was -5 (of a possible 9 if all comparisons were significant in the same direction), indicating moderate certainty of results.

The Normalized Enrichment Scores (NES) by GSEA for pathways enriched by Fisher’s method are presented in **Figure 2G**. Strong heterogeneity is observed across comparisons, but some patterns can be detected: the two studies profiling expression in mouse blood or human lymphoid-derived cells cluster together, while the seven from brain tissue comprise the other cluster. A group of immune pathways show strong enrichment in resistant blood samples, but weak, opposite enrichment across brain samples from responding mice. Other clusters of pathways show heterogeneous patterns within the set of comparisons profiling brain tissue, and none are enriched in the same direction across all comparisons, as evidenced by the distribution of vote sums.

Next, we focused on pathways that were identified as consistently enriched across comparisons with the Max-P method. Meta-analysis statistics for pathways with q<0.05 are presented in **Figure 3A**, and a network diagram showing similarity between these gene sets is presented in **Figure 3B** (a heatmap showing overlap between gene sets is provided in **Supplementary Figure 1**). Signal Transduction (a top-level Reactome pathway consisting of over 2000 genes) was the most consistently enriched pathway with q<0.001 by Max-P and a vote sum of +3, indicating somewhat consistent enrichment in good responders. 10 immune pathways were identified as consistently enriched, including five toll-like receptor (TLR) pathways and two MyD88 cascade pathways; these seven pathways are highly overlapping, sharing over 95% of genes. Additionally, the NF-kappa B signaling pathway, C-type lectin receptors (CLRs), Downstream signaling of B Cell Receptor (BCR), and the top-level Immune System pathway were identified by meta-analysis. These were slightly enriched in good responders by total vote, except for the CLRs pathway with a vote sum of -1 (significantly enriched in non-responders in patient-derived LCL’s).

**Figure 3.**
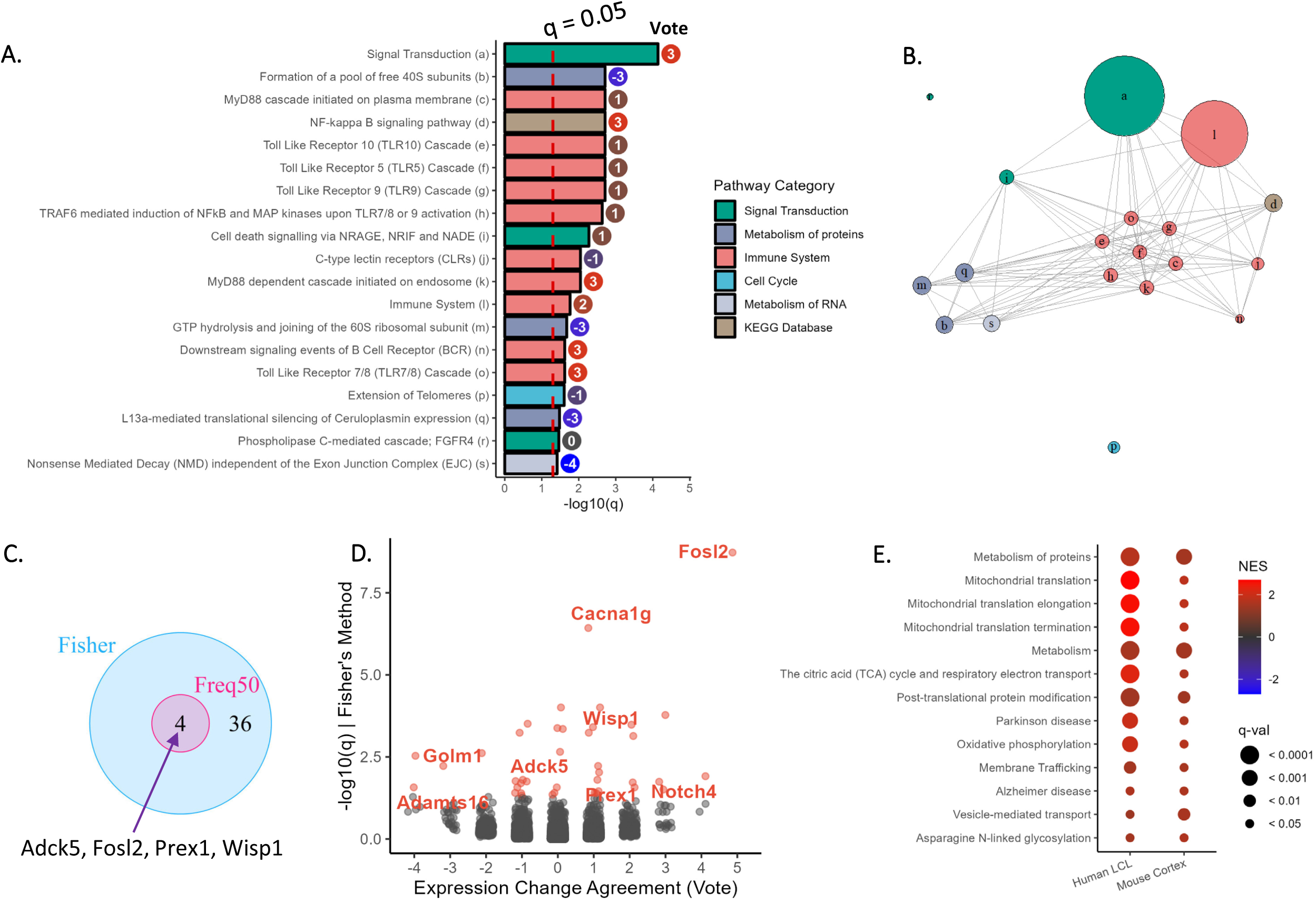
(A) q-value by Max-P and Vote Sums for pathways identified as enriched across the nine comparisons of responders vs. non-responders (q < 0.05 by Max-P). **(B)** Network diagram showing similarity between pathways, with labels provided in ***3A***. Pathways sharing more genes are plotted closer together. **(C)** Meta-analysis results for genewise differential expression in eight mouse comparisons. No genes were identified as consistently differentially expressed by Max-P. **(D)** Volcano plot of mouse differential expression meta-analysis results. **(E)** 13 pathways enriched in responders vs. non-responders in two comparisons of samples not treated with fluoxetine (q < 0.05 in each).

Pathways related to metabolism of proteins or RNA were most consistently enriched in poor responders, including the substantially overlapping pathways related to 40S and 60S ribosomal subunits (both with vote sums of -3), L13a-mediated translational silencing of Ceruloplasmin expression (-3), and Nonsense Mediated Decay independent of the Exon Junction Complex (-4). These were enriched in non-responders in both comparisons involving blood-derived tissue, as well as some comparisons in brain tissue. Additionally, Extension of Telomeres (q=0.02, with negligible overlap with the other enriched pathways shown in **Figure 3B**) was enriched by Max-P with a vote sum of -1.

Meta-analysis was conducted for differential expression at both the gene and pathway level for the eight comparisons within mice. More pathways were identified as consistently differentially expressed by Max-P (27 with only mouse studies vs. 19 including the patient cohort), although many of the uniquely identified pathways also involved TLR cascades (**Supplementary Figure 2**). Individual gene meta-analysis proved less consistent, with only 40 genes identified as differentially expressed in *any* comparison by Fisher’s method, four differentially expressed in over half of the comparisons, and none identified as consistent by the Max-P method (**Figure 3C**). A meta-volcano plot of these results is presented in **Figure 3D**. *Fosl2* is both highly enriched by Fisher’s method (q<0.001) and consistently overexpressed in good responders (vote sum of +5). *Notch4* (+4), *Golm1* (-4), and *Adamts* (-4) are the other genes that are most consistently differentially expressed across comparisons in mice.

We conducted two sensitivity analyses successively removing individual comparisons from the meta-analysis. The previously presented meta-analysis across mouse studies demonstrated the effects of removing the single patient data set; we then conducted a second analysis synthesizing the seven mouse comparisons in brain tissues only. Overlap between pathways identified by Fisher’s and Max-P methods can be seen in **Supplementary Figure 2**. As individual data sets were removed, pathways identified by Fisher’s method decreased, as this method identifies whether a pathway is enriched in *any* of the studies. By Max-P, only two pathways were identified in the full set but not in the subsets, while ten additional pathways were identified when the patient comparison was removed (as previously discussed). Meta-analysis of the seven brain comparisons did not identify any additional pathways by Max-P: thus, we can conclude that the Max-P meta-analysis of responders vs. non-responders was only slightly sensitive to the inclusion of gene expression profiling in blood samples, as all pathways identified as consistently enriched in responders or non-responders in brain were also enriched in blood (although not necessarily in the same direction).

Finally, we compared results from two studies that profiled gene expression in responders and non-responders that had not previously received fluoxetine to identify potential predictive signatures: one in patient-derived LCL’s and the other in mouse cortex samples (mouse response was inferred based on overall response measured for multiple mice from each strain)(45). 13 pathways were identified as enriched by GSEA with q<0.05 in both comparisons, and all were enriched in good responders (**Figure 3E**). These included pathways related to metabolism and mitochondrial translation, in addition to gene signatures of Alzheimer’s disease and Parkinson’s disease.

### Treatment Signatures

The same meta-analyses were applied on 55 comparisons of treated vs. control samples from 20 data sets, to synthesize gene expression changes due to treatment (**Figure 4A**). First, each data set was systematically reanalyzed, again providing widely varying results (**Figure 4B**): after false discovery correction, the number of differentially expressed genes ranged from zero (in 19 comparisons) to 8663, and the number of significantly enriched pathways ranged from zero (in 7 comparisons) to 971. Fisher’s meta-analysis identified 737 pathways as enriched in at least one comparison; of these, 31 were enriched with nominal p<0.05 in more than half of the comparisons, and 10 were identified as consistently enriched by the Max-P method (**Figure 4C**).

**Figure 4.**
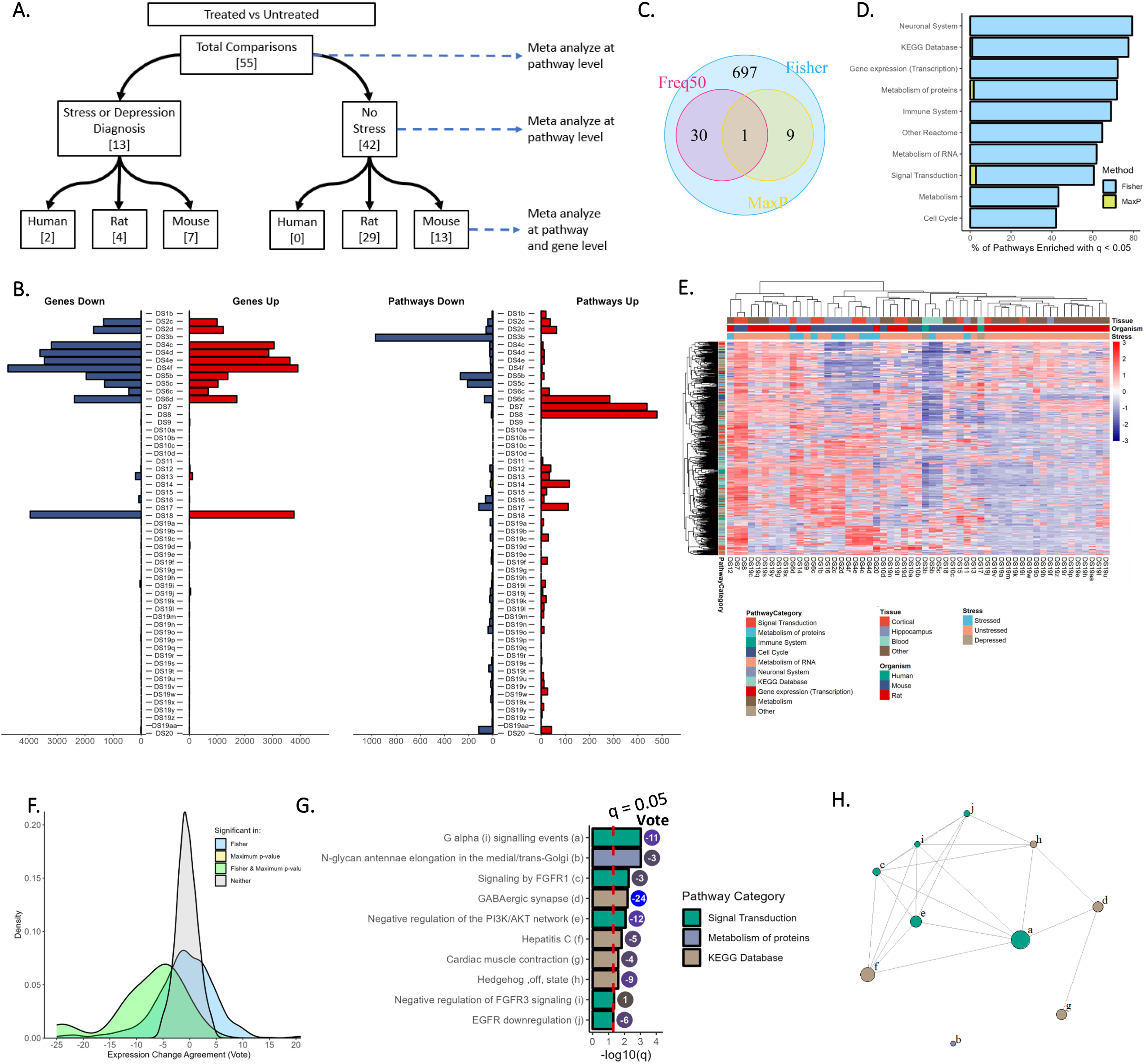
(A) Individual comparisons used in meta-analysis for gene expression signatures of fluoxetine treatment effects. **(B)** Number of genes differentially expressed (left) and pathways enriched (right) in each of the nine comparisons of samples from fluoxetine treated vs. control samples (q < 0.05). “Up” indicates increased expression with fluoxetine. **(C)** Pathways identified as significantly enriched (q < 0.05) across the 55 comparisons by the three meta-analysis methods. **(D)** Percentage of pathways significantly affected by fluoxetine treatment from each of the Reactome categories (“Top Level”) or KEGG Database. **(E)** Heatmap of Normalized Enrichment Score (NES) across comparisons for pathways identified as enriched in any comparison (Fisher’s q < 0.05). Positive NES indicates increased expression with fluoxetine treatment. **(F)** Density plot showing distribution of Vote Sums across pathways, colored by meta-analysis result. **(G)** q-value by Max-P, and Vote Sums for pathways identified as significant across the 55 comparisons of treated vs. untreated (q < 0.05 by Max-P). **(H)** Network diagram showing similarity between pathways, with labels provided in ***4G***. Pathways sharing more genes are plotted closer together.

The pathways identified as enriched in any of the 55 comparisons (65% of all pathways analyzed) are represented by all pathway categories, but most frequently components of Neuronal System or the KEGG database (**Figure 4D**). Heterogeneous enrichment patterns are again observed across studies (**Figure 4E**). Tissue type again played a role, as three of the four comparisons from blood clustered tightly together, and most comparisons from hippocampus and cortex clustered together. Comparisons from the same study frequently clustered together as well, evidenced by the data set ID’s.

Overall, pathways identified only by Fisher’s method have a wider distribution of vote sums (but still highly concentrated between +6 and -6, of a possible 55 for perfect agreement) than those that are not significantly enriched, while those identified by Max-P are skewed toward the left indicating downregulation by treatment (**Figure 4F**). The 10 pathways enriched by Max-P (**Figure 4G**) mainly come from the Signal Transduction category of Reactome or the KEGG Database. GABAergic synapse is most consistently decreased by treatment with a vote sum of -24 (q=0.006). G alpha (i) signaling events are identified as most enriched by q-value and somewhat consistently decreased by treatment (q=0.001, -11). There was not substantial overlap between genes within enriched pathways (**Figure 4H** and **Supplementary Figure 3**).

Subset meta-analysis was conducted for 13 comparisons from 10 data sets profiling fluoxetine treatment of stressed mice, rats, or human MDD patients. 731 pathways were identified as enriched in at least one comparison, and 101 were identified as consistently enriched by Max-P (**Figure 5A**). Greater enrichment is observed in immune pathways in this subset analysis as compared with the full analysis (**Figure 5B**). Generally low agreement between studies is observed, with vote sums ranging from -2 to +2 for 98.7% of pathways not identified as enriched by meta-analysis (**Figure 5C**). 21% of pathways identified by both Fisher’s and the Max-P method had vote sums outside this range.

**Figure 5.**
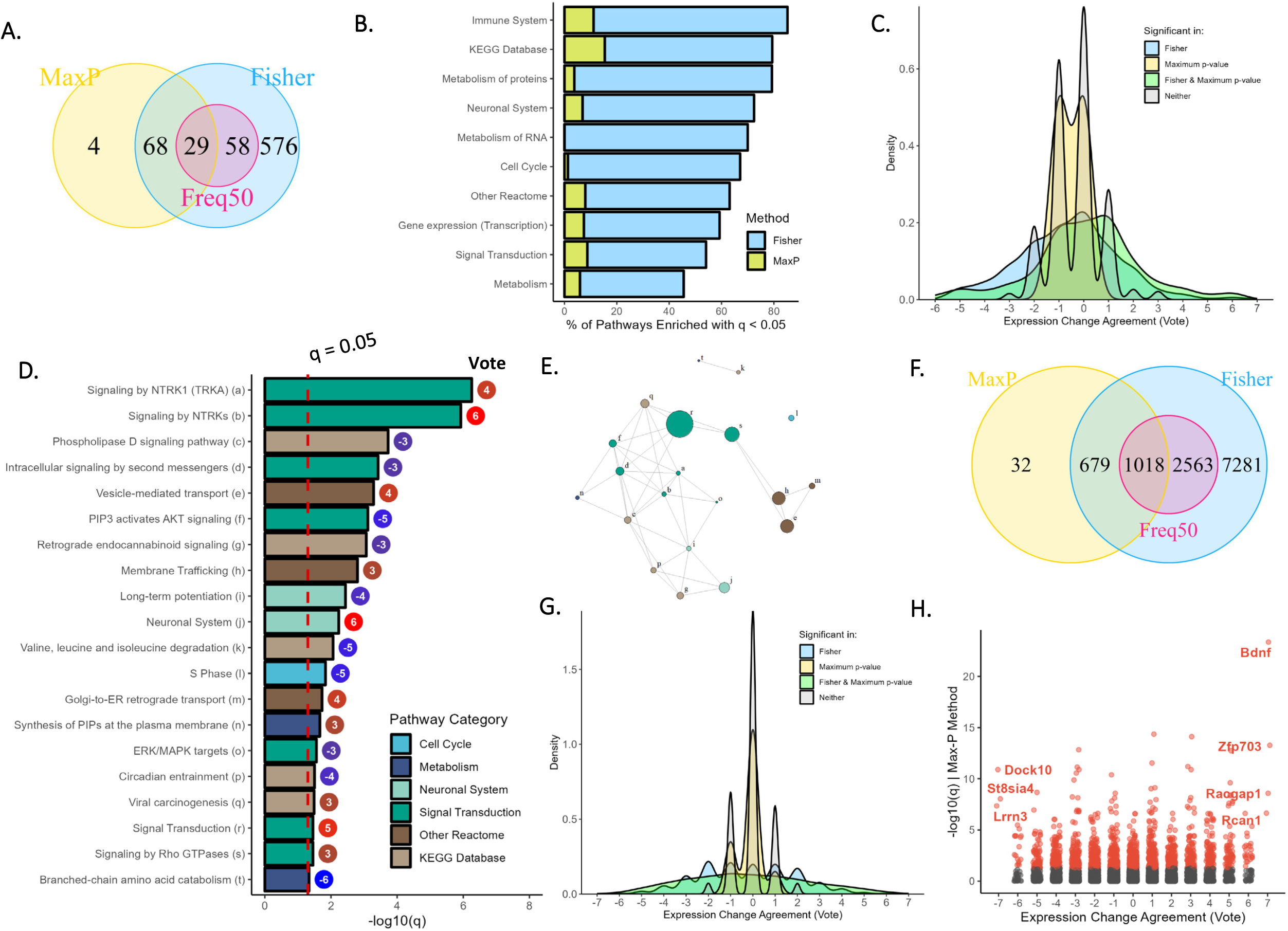
(A) Pathways identified as significantly enriched (q < 0.05) across the 13 comparisons of treatment effects in MDD patients or stressed rodent models. **(B)** Percentage of pathways significantly affected by fluoxetine treatment in MDD patients or stressed rodent models from each of the Reactome categories (“Top Level”) or KEGG Database. **(C)** Density plot showing distribution of Vote Sums across pathways, colored by meta-analysis result. Positive scores indicate increased expression with fluoxetine. **(D)** q-value by Max-P and Vote Sums for pathways identified as significant across the 13 comparisons in MDD patients or stressed rodents (q < 0.05 by Max-P), subset to include only those with absolute Vote Sum ≥ 3. **(E)** Network diagram showing similarity between pathways, with labels provided in ***5D***. **(F)** Genes identified as differentially expressed (q < 0.05) across the seven comparisons of treatment effects in stressed mice. **(G)** Density plot showing distribution of Vote Sums across genes, colored by meta-analysis result. **(H)** Volcano plot of meta-analysis results for differential expression of fluoxetine vs. control in stressed mice. Genes with absolute vote sum of 7 are labeled.

Of the pathways identified by Max-P, we highlighted those with absolute vote sums of 3 or greater in **Figure 5D**, with gene set similarity presented in **Figure 5E** (all 101 pathways are presented in **Supplementary Figure 4**). Most significantly enriched are two NTRK signaling pathways, both with q<0.001 and vote sums of +4 and +6 respectively (of 13 possible), indicating moderate certainty. Other signal transduction pathways show a decrease with treatment, including Intracellular signaling by second messengers (q<0.001, vote sum of -3) and PIP3 activates AKT signaling (q=0.001, -5). The top-level Neuronal System gene set is significantly upregulated with treatment (q=0.005, +6); conversely, the component Long-term Potentiation pathway is downregulated (q=0.003, -4). A network graph showing overlap between all pathways identified by Max-P indicates groups of connected pathways from the KEGG Database and Reactome Signal Transduction, in addition to smaller groups of other pathways (**Figure 5E**).

A summary of all within-species meta-analysis results (bottom level of **Figure 4A**) is presented in **Supplementary Figure 5**. Profiling treatment effects in stressed mice identified the greatest number of differentially expressed genes, with 11,541 identified by Fisher’s method and 1729 identified as consistently differentially expressed by Max-P (**Figure 5F**). Genes identified by both Fisher’s method and Max-P had a wider distribution of vote sums, with 39.8% with absolute value of 3 or greater (**Figure 5G**). Seven genes were significantly affected by treatment in the same direction across all seven comparisons: *Bdnf, Zfp703, Raogap1, Rcan1, Dock10, St8sia4,* and *Lrrn3* (**Figure 5H**). Of these, Brain-derived neurotrophic factor (*Bdnf*) was identified as most significantly affected by Max-P (q<<0.001).

Overall, low agreement is observed between the meta-analysis of stressed rodents or MDD patients and the meta-analysis of unstressed rodents. No pathways overlap by Max-P, although 425 pathways overlapped by Fisher’s method (**Supplementary Figure 6**). We conducted a sensitivity analysis removing DS19 (profiling across 27 brain regions), as this was over half of the comparisons included in the synthesis, some of which were reported as minimally affected by fluoxetine(59). We expected our Max-P results to be sensitive to comparisons in less-affected brain regions--conversely, we saw that fewer pathways were consistently enriched upon removal of this study (**Supplementary Figure 7**). Considering the gene level, 149 genes were consistently differentially expressed in the original meta-analysis, vs. none when DS19 was removed. Thus, we can conclude that meta-analysis of treatment effects was sensitive to DS19. As seen in **Figure 4E**, comparisons across the 27 tissue types in DS19 showed largely consistent patterns, likely contributing to the observed result.

Finally, we conducted separate analyses and meta-analyses within responder and non-responder groups across studies that provided response information, to identify gene expression changes that may indicate response to treatment. The Max-P method identified 22 pathways that were consistently changed across both groups, 86 pathways that were specific to responders, and 29 that were specific to non-responders (**Figure 6A**). Of the pathways that consistently changed with treatment in responders, we focused on the 17 that were consistently *unchanged* in non-responders (**Figure 6B**, q>0.05 in non-responders by Max-P and Fisher’s method). This included eight pathways from the KEGG database, five signal transduction pathways, two immune pathways, and two others. However, vote sums ranged from -3 to +3, indicating weak certainty of evidence. Conversely, only the estrogen signaling pathway was consistently changed in non-responders but consistently unchanged in responders based on the same criteria.

**Figure 6.**
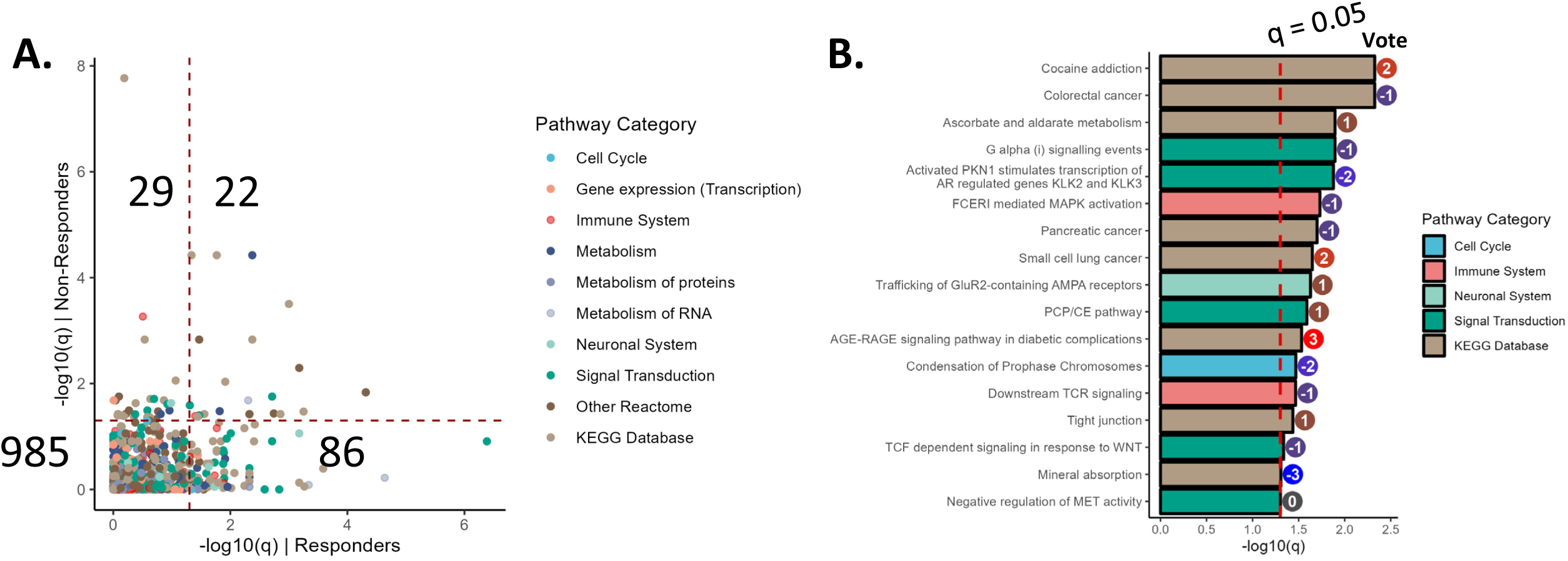
(A) Comparison of pathway enrichment by treatment for responders and non-responders, by Max-P. Dotted lines indicate q = 0.05, and counts of pathways within each quadrant are provided. **(B)** q-value by Max-P and Vote Sums for pathways identified as significant across the eight comparisons of treatment vs. control in responders (q < 0.05 by Max-P), but consistently *not* significant across eight comparisons of treatment vs. control for non-responders (q > 0.05 by both Max-P and Fisher’s method).

## DISCUSSION

Across nine independent comparisons of fluoxetine responders vs. non-responders from six gene expression data sets, 19 pathways were identified as consistently enriched by the Max-P meta-analysis method. This was surprising, as these studies profiled different tissue types and employed varying stress models in mice--in addition to one which did not induce stress, and another using samples from MDD patients. This also demonstrated the ability of pathway methods vs. individual gene meta-analysis in identifying consistency: among the eight comparisons in mice, zero genes were identified with consistent differences between responders and non-responders by Max-P, and only four were identified as differentially expressed in greater than half of comparisons. Less surprisingly, over three hundred pathways were identified as enriched in at least one comparison by Fisher’s method, demonstrating the expected heterogeneity in behavioral response markers between studies. Some of this heterogeneity can be explained by tissue type, as comparisons from similar tissues clustered nearest each other (**Figure 2G**). Thus, we also conducted separate meta-analysis excluding studies profiling expression in blood, but we saw only a slight effect on the meta-analysis result by Max-P, indicating that pathways enriched in brain tissue were generally enriched in blood as well, but sometimes in the opposite direction.

Immune pathways were well represented among the 19 pathways that were consistently different between responders and non-responders across organisms and tissue types. Many studies have implicated immune and inflammatory processes in the development of depression and antidepressant response(63–70). For example, Wittenberg *et al.* demonstrated that anti-IL-6 and anti-IL-12/23 antibodies improve depressive symptoms in patients with inflammatory or oncological disorders(71). Of the immune pathways identified, TLR-related pathways were the most enriched in our meta-analysis. TLRs are present in microglia and other glial cells, and their activation leads to the release of cytokines and other inflammatory responses. Rodents exposed to stress or glucocorticoid administration have been shown to exhibit increased levels of TLR2 and TLR4, while blockade of TLR2 and TLR4 has been shown to prevent neuroinflammatory response(70,72–75). Additionally, one study included in our meta-analysis reported the involvement of inflammation via the TLR pathway in depression pathogenesis and investigated the use of acupuncture in relieving this inflammation(54). The consistent enrichment of the NF-κB signaling pathway upon meta-analysis is connected to this pattern, as activity of this pathway has been shown in rodents to mediate depressive-like behavior and increase release of pro-inflammatory cytokines in the microglia(76).

However, the direction of immune enrichment across the meta-analyzed studies was not consistent: while immune pathways were strongly upregulated in resistant samples in the two studies profiling gene expression in blood (including one study in human patients), these pathways were weakly upregulated in responding mice in studies profiling brain tissues. A recent systematic review has noted that inflammatory markers are only weakly correlated between peripheral and cerebrospinal fluid(77). Carillo-Roa *et al.* identified 85 differentially expressed genes (q<0.05) in the blood of responders vs. non-responders in mice treated with paroxetine, but similar differential expression was not observed in mouse prefrontal cortex; importantly, they demonstrated that these 85 genes could correctly predict response with 81% accuracy in humans treated with duloxetine or escitalopram, demonstrating that transcripts found in blood may be an accessible, objective diagnostic marker even if expression patterns are not shared in brain(78). Additionally, Le-Niculescu *et al.* have identified biomarkers for mood disorders that are present in both brain and blood, providing additional evidence for the value of these markers(23).

Protein and RNA metabolism pathways were among the most consistently enriched in samples resistant to fluoxetine treatment across organisms and tissue types. Specifically, two pathways involving the 40S and 60S ribosomal subunits were enriched; a few studies have previously identified ribosomal proteins and pathways as implicated in response to antidepressants(79,80). Additionally, Zhou *et al.* have reported evidence that ribosomes regulate gene expression involved in immune response(81).

Tissue heterogeneity again played a role when considering gene expression changes due to fluoxetine treatment, with blood-derived tissues showing distinct overall effects from those in the brain (**Figure 4E**). However, considering the 55 widely varying comparisons of fluoxetine treatment vs. control, ten pathways were identified as consistently differentially expressed with the Max-P method (**Figure 4G**). Of these, the GABAergic synapse was most downregulated by fluoxetine. GABAergic neurons have been implicated in depression and antidepressant response, and positive modulators of the GABA_A_ receptor have been approved by the FDA for postpartum depression(82–85). Additionally, we identified BDNF as most consistently upregulated with treatment in stressed mice (q<<0.001 by Max-P, upregulated with p<0.05 in all seven studies); Tanaka *et al.* observed that BDNF inhibits the GABA_A_ synaptic response in rat hippocampus(86). Evidence from rodent studies has also suggested bidirectional connections between BDNF expression and inflammation, as inflammation via lipopolysaccharide treatment was shown to increase BDNF secretion in the microglia, while application of BDNF in spinal cord injury has been shown to decrease microglial density(87–89). Other studies have demonstrated similar connections, indicating that elevation of BDNF by fluoxetine may participate in amelioration of depression symptoms by both GABAergic and anti-inflammatory effects(90–92).

When considering only stressed models, many more pathways (101) were identified as consistently affected by fluoxetine treatment (**Figure 5A**). Further separation into treatment effects of responders and non-responders demonstrated a greater number of pathways consistently affected by treatment in responders than non-responders (**Figure 6A**). Effects were again observed in signal transduction and immune pathways, while pathways related to cancer and addiction were also identified. Antidepressants have recently been demonstrated to inhibit liver and lung cancer through the mTOR pathway, although other evidence has indicated associations between antidepressant use and increased cancer incidence(93–95).

While the PRISMA guidelines are more commonly applied for systematic review of clinical studies, we felt that they provided a strong framework for this work. Systematic identification and re-analysis of each data set allowed us to calculate consistent metrics for differential expression, rather than relying on lists of statistics provided by individual study authors. We used broad inclusion criteria for organism and tissue to determine whether consistent changes were observable across heterogeneous studies; the use of Fisher’s meta-analysis method, subgroup analyses, and sensitivity analyses then let us explore the expected heterogeneity. Reporting bias is challenging to assess in gene expression analysis, where thousands of statistics are generated for each study. Yousefi *et al*. have assessed risk of reporting bias when classification algorithms are applied to gene expression data(96), but we are not aware of methods to detect reporting bias of the gene expression data itself; this will be a valuable tool as systematic reviews of gene expression studies become more prevalent.

This study did include multiple limitations that should be addressed to better understand antidepressant effectiveness. Only two studies provided gene expression data for responders and non-responders prior to treatment, so it is not possible to derive strong inferences for predictive markers of fluoxetine response; even large-scale patient studies have struggled to identify general predictive markers to this point(18,97). Only two patient cohorts were included in our meta-analyses: while other studies have been conducted in patients, they tend to focus on a small number of biomarkers or otherwise do not submit full expression data to GEO(18). Most data sets profiled in our meta-analyses contained sample sizes of five or fewer per group, although some samples were pooled across multiple rodents. Additionally, this systematic review was not prospectively registered, and the reviewers did not work independently (all exclusion decisions are documented in **Supplementary Table 1**).

The included studies showed substantial bias based on sex, with 17 of 18 rodent studies profiling males, while the two patient studies included majority or exclusively female participants. Results of Max-P meta-analysis provide some evidence for conserved response and treatment signatures, but it is not possible to identify sex-dependent biological signatures with these data. Previous work has shown both consistency and divergence between males and females regarding clinical response, molecular signatures, and adverse effects from antidepressants, and this remains an important area of study(43,98–101).

Considering the meta-analysis methods, reliance on the Max-P method means that we are particularly sensitive to a single aberrant study resulting in high p-values due to data quality or an unconsidered factor, which may not have been apparent during screening. For this reason, all code and results are provided in our registered repository for further inspection and analysis, including individual data set analyses, as well as other meta-analyses by Fisher’s method (which is more sensitive to studies that may have extremely low p-values) and the Freq-50 method (which may be most robust but does not test a specific hypothesis).

Our meta-analyses have emphasized some known pathways in antidepressant response and unearthed a few new routes of potential investigation, but a true understanding of antidepressant response will require additional large-cohort studies focusing on transcriptomic data. However, recent studies of antidepressant response in over 100 patients have resulted in few, if any, individual biomarkers of response, supporting the consensus that there is not a single genetic signature of response, but a variety of contributing ‘omic and environmental factors(18,97). Emerging technologies and approaches have begun to allow us to understand genomics and transcriptomics at a deeper level, including single-cell profiling, alternative splicing, and epigenetic factors(59,102–104). And as technology improves our ability to quantify protein and metabolite levels, it will be valuable to incorporate these data, as this will bring us even closer to the true biology underpinning these complex disorders.

## Acknowledgements

The authors gratefully acknowledge John Nurnberger Jr., M.D., Ph.D. (Indiana University School of Medicine); C. Andrew Class, M.D. (Ascension St. Vincent); and Marcos Oliveira, Ph.D. (Butler University) for helpful suggestions and conversations. This work was supported by the American Association of Colleges of Pharmacy (AACP New Investigator Award to CAC), the Butler University Holcomb Awards Committee, and departmental funds. Funders were not directly involved in the execution of this project.

## Competing Interests

The authors declare no conflict of interest.

## Code Availability

All analyses completed in this work are included at https://github.com/DavidGCooper/FLX-MetaDE with DOI 10.5281/zenodo.10668845(105).

---

**Supplementary Figure 1.**
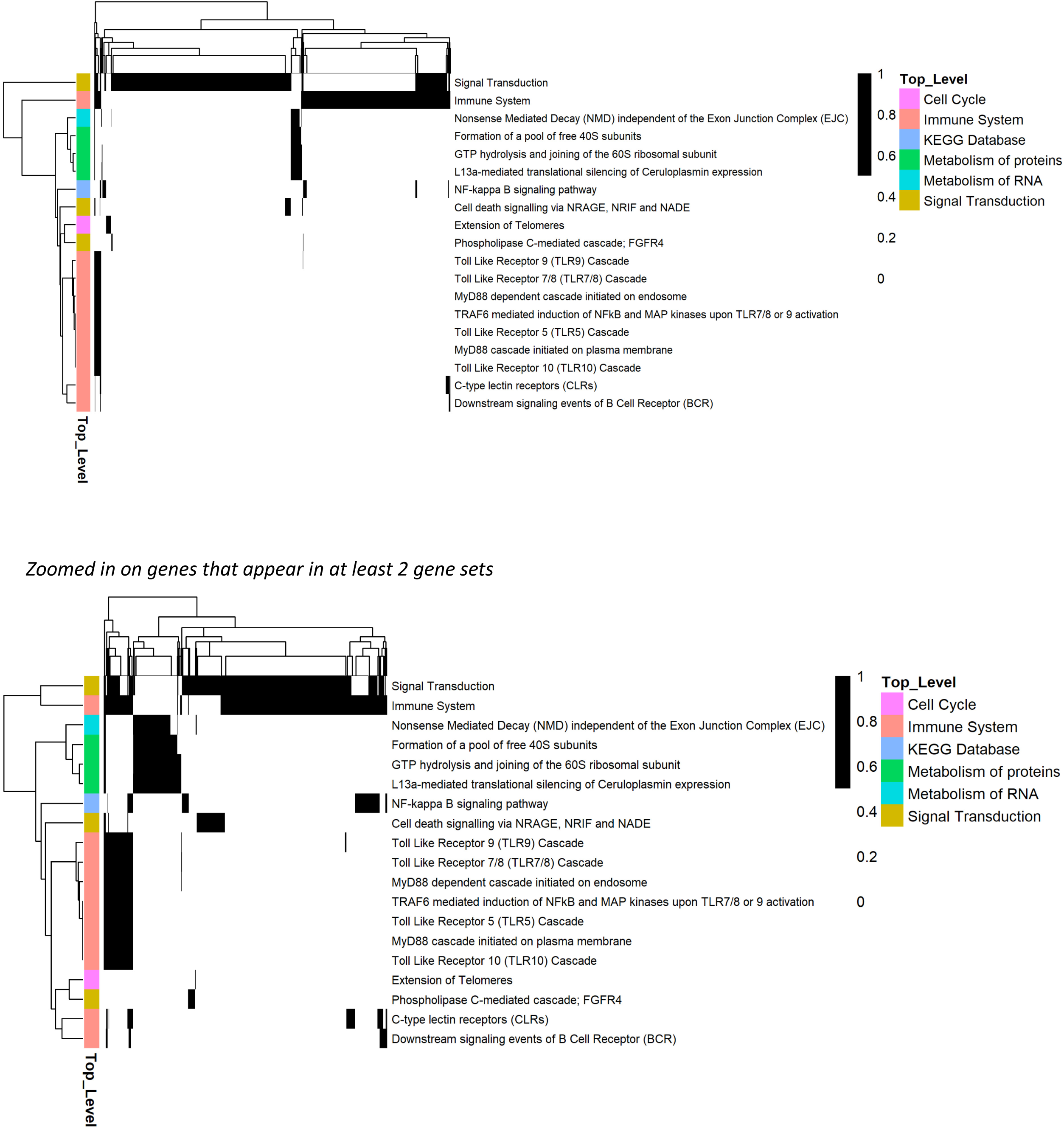
Overlap of genes included in pathways identified as significant across the nine comparisons of responders vs. non-responders (q < 0.05 by Max-P). (*Top*) All included genes. (*Bottom*) Subset of genes appearing in at least two of the pathways.

**Supplementary Figure 2.**
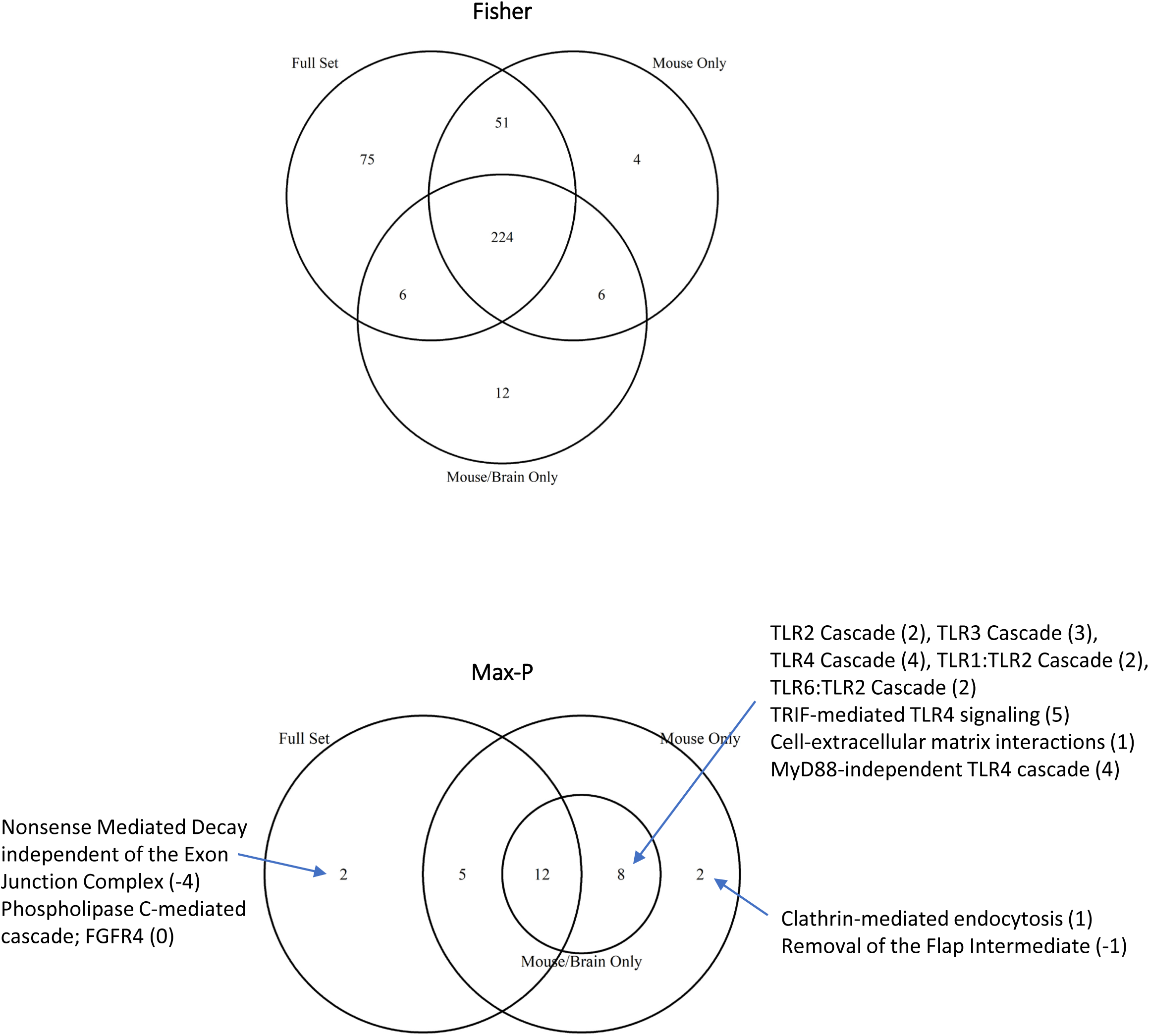
Overlap of pathways identified for responders vs. non-responders for all nine comparisons, with two sensitivity analyses (meta-analysis with human comparison removed, and meta-analysis with blood comparisons removed). Vote sums are indicated in parentheses.

**Supplementary Figure 3.**
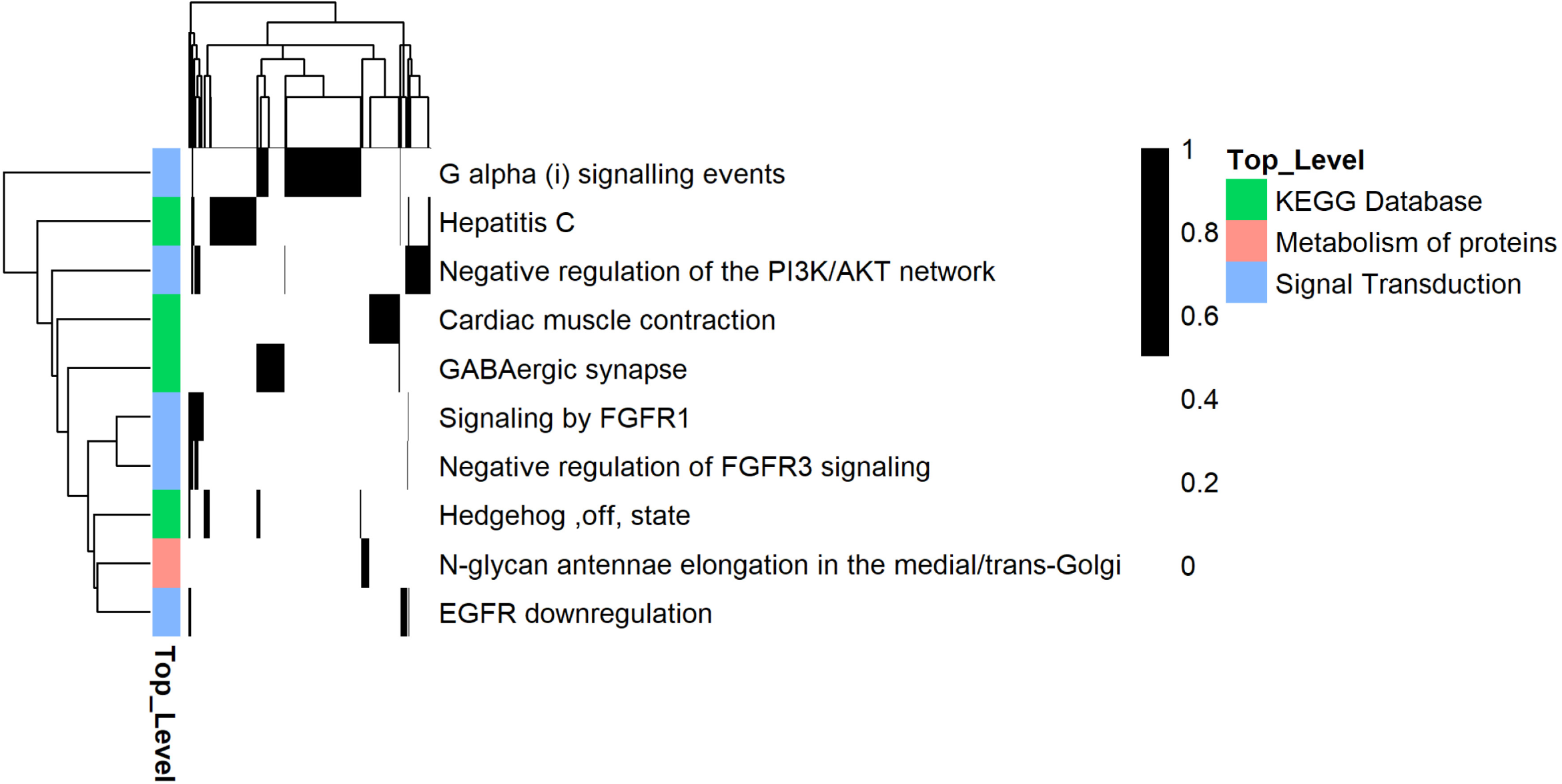
Overlap of genes included in pathways identified as significant across all comparisons of fluoxetine treated vs. untreated samples (q < 0.05 by Max-P).

**Supplementary Figure 4.**
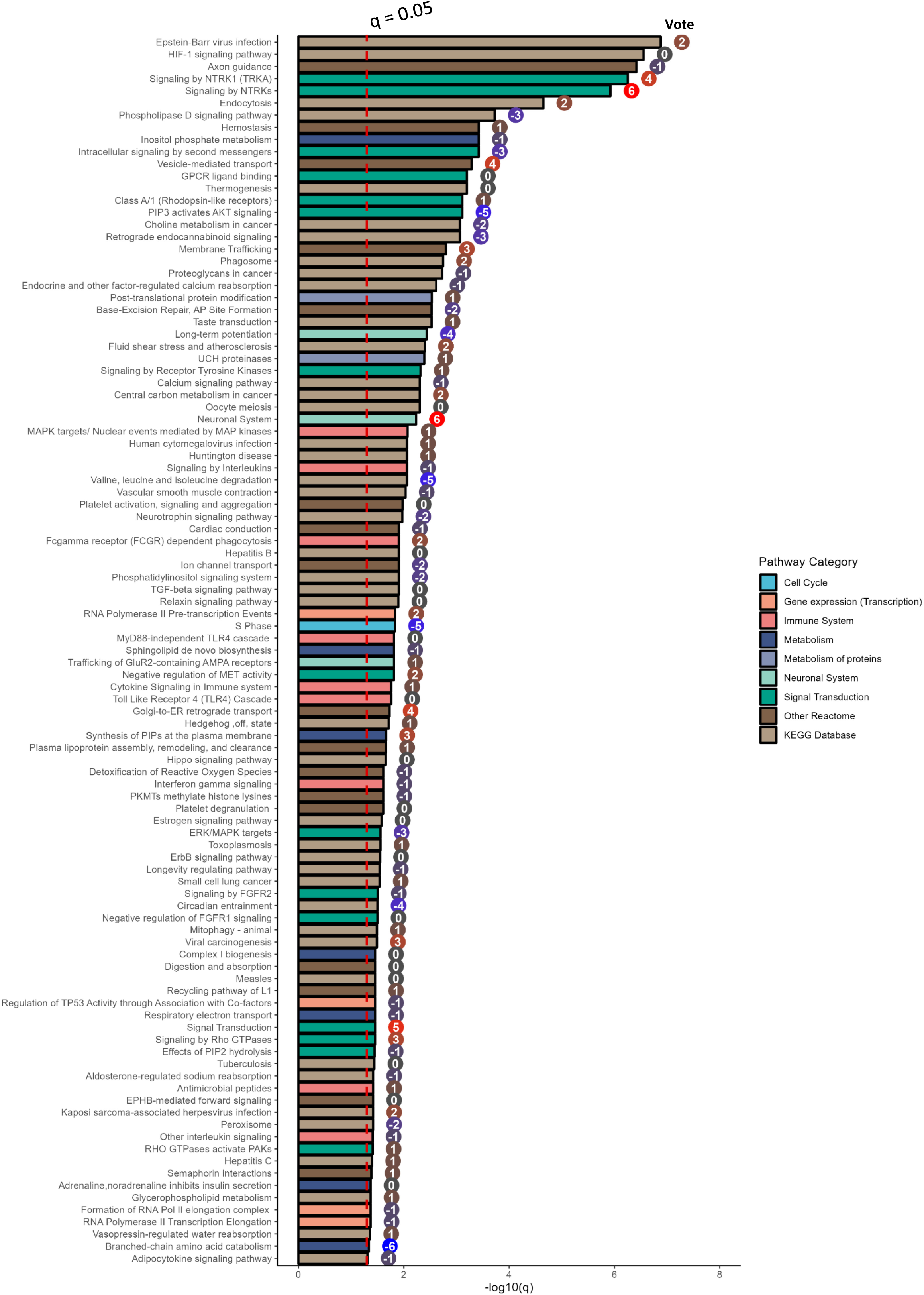
q-value by Max-P and Vote Sums for all pathways identified as significant across the 13 comparisons in MDD patients or stressed rodents (q < 0.05 by Max-P).

**Supplementary Figure 5.**
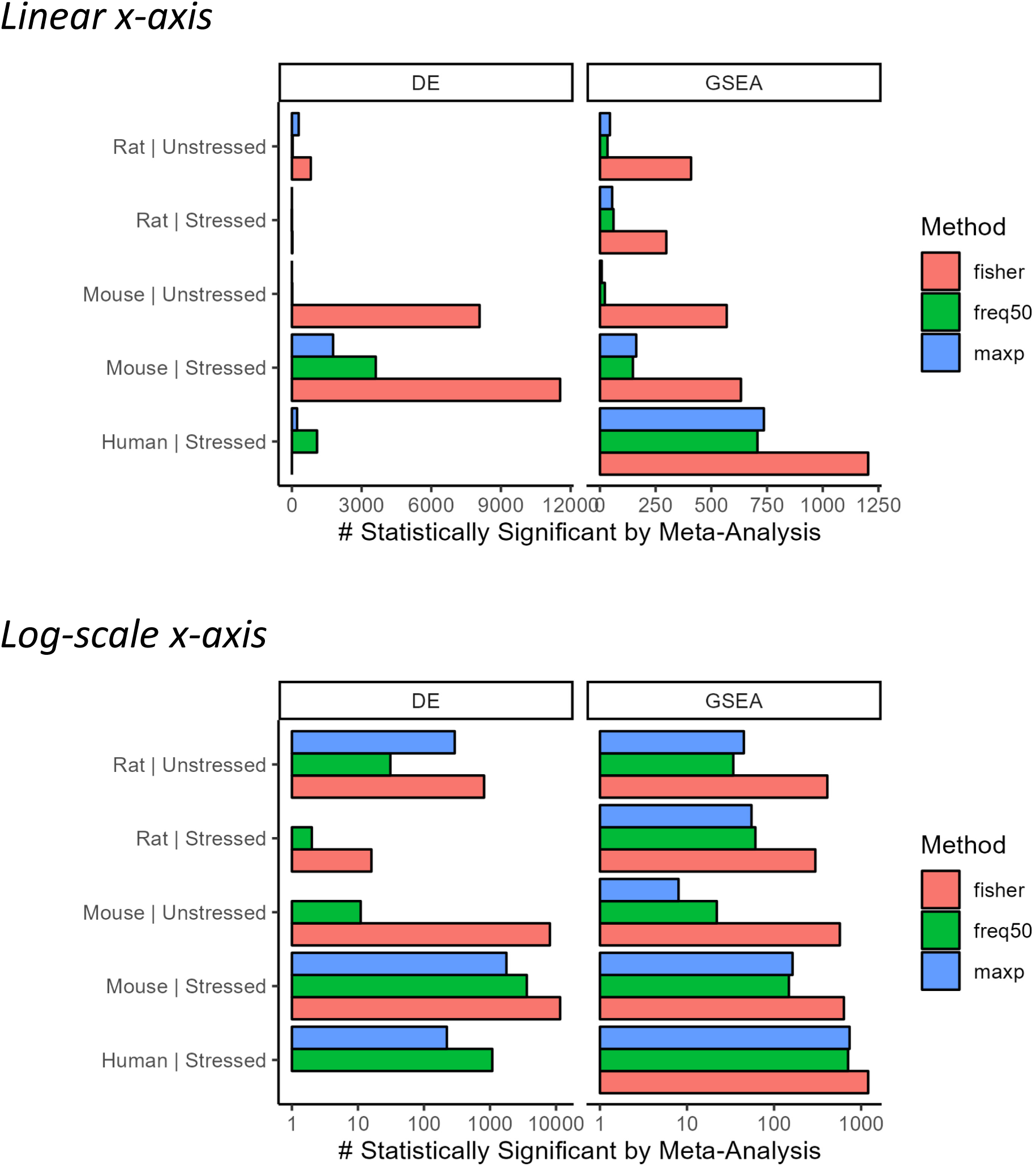
Number of genes (“DE”) or pathways (“GSEA”) identified as statistically significant in each meta-analysis, with q < 0.05. Same data are presented in linear (*top*) and log10 scale (*bottom*).

**Supplementary Figure 6.**
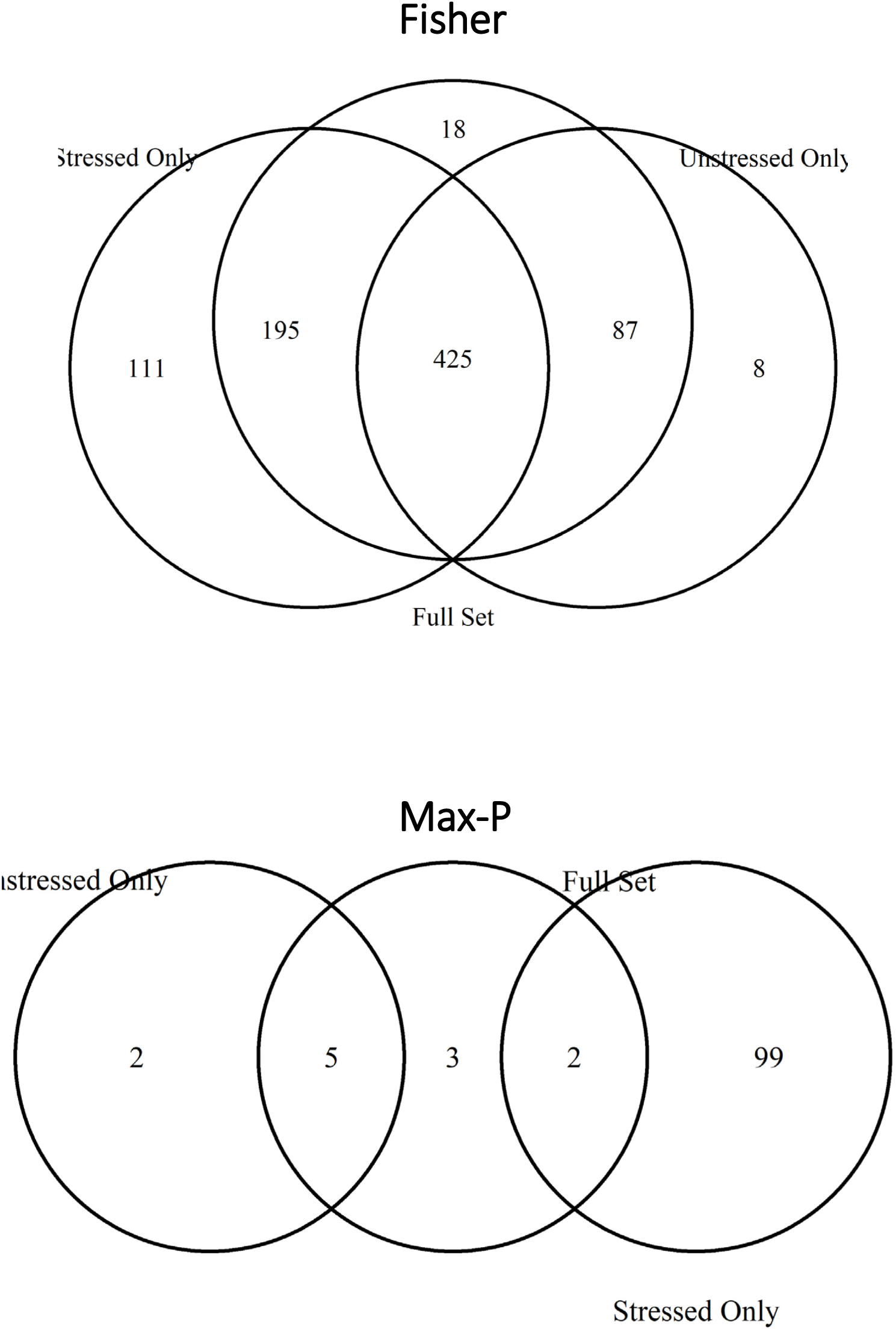
Number of pathways identified as statistically significant for the full meta-analysis of treated vs. untreated samples, as well as subset analyses of stressed rodents or depressed patients, and unstressed rodents. Results of Fisher’s meta-analysis (*top*) and Max-P (*bottom*).

**Supplementary Figure 7.**
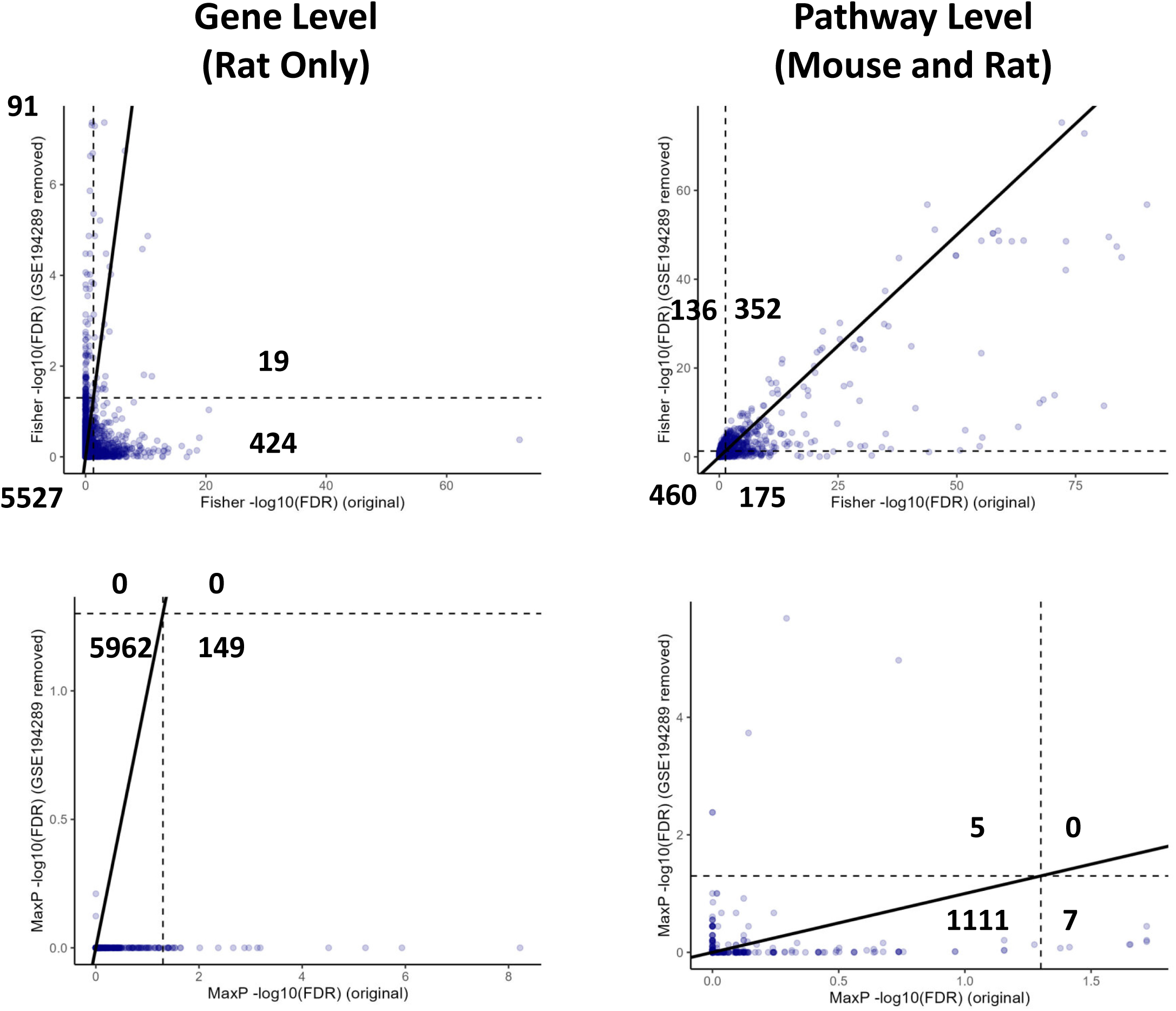
Sensitivity analysis removing DS19 (profiling of 27 brain regions by Rayan *et al.*(59)). Gene (*left*) and pathway-level (*right*) analyses are presented. Dashed lines indicate q=0.05, diagonal line indicates y=x. Number in each quadrant indicates number of genes or pathways in that quadrant. Results of Fisher’s meta-analysis (*top*) and Max-P (*bottom*).

## Notes

https://zenodo.org/doi/10.5281/zenodo.10668845

### Competing Interest Statement

The authors have declared no competing interest.

### Summary of Updates

The discussion section was revised to better place this work in the context of existing and recent studies.

